# Mammalian stringent-like response mediated by the cytosolic NADPH phosphatase MESH1

**DOI:** 10.1101/325266

**Authors:** Chien-Kuang Cornelia Ding, Joshua Rose, Jianli Wu, Tianai Sun, Kai-Yuan Chen, po-Han Chen, Emily Xu, Sarah Tian, Jadesola Akinwuntan, Ziqiang Guan, Pei Zhou, Jen-Tsan Chi

## Abstract

Nutrient deprivation triggers stringent response in bacteria, allowing rapid reallocation of resources from proliferation toward stress survival. Critical to this process is the accumulation/degradation of (p)ppGpp regulated by the RelA/SpoT homologues. While mammalian genomes encode MESH1, a homologue of the bacterial (p)ppGpp hydrolase SpoT, neither (p)ppGpp nor its synthetase has been identified in mammalian cells. Therefore, the function of MESH1 remains a mystery. Here, we report that human MESH1 is an efficient cytosolic NADPH phosphatase, an unexpected enzymatic activity that is captured by the crystal structure of the MESH1-NADPH complex. MESH1 depletion promotes cell survival under ferroptosis-inducing conditions by sustaining the level of NADPH, an effect that is reversed by the simultaneous depletion of the cytosolic NAD(H) kinase, NADK, but not its mitochondrial counterpart NADK2. Importantly, MESH1 depletion also triggers extensive transcriptional changes that are distinct from the canonical integrated stress response but resemble the bacterial stringent response, implicating MESH1 in a previously uncharacterized stress response in mammalian cells.

---

Stringent response is the main strategy for bacteria to cope with fluctuating nutrient supplies and metabolic stresses ^1,2^. During this process, alterations in transcriptional and metabolic profiles rapidly redirect energy from cell proliferation toward stress survival by reduction of biosynthesis, conservation of ATP, and blockage of GTP production ^3^. The stringent response is triggered by the accumulation of the bacterial “alarmone” (p)ppGpp (guanosine tetra-or penta-phosphate, shortened as ppGpp below) through the regulation of ppGpp synthetases and hydrolases in the RelA/SpoT Homologue family ^2^. Recent studies suggest that the stringent response may also function in metazoans, as metazoan genomes encode a homologue of bacterial SpoT, MESH1 (<u>*Me*</u>tazoan <u>*S*</u>poT <u>*H*</u>omolog 1) that hydrolyzes ppGpp *in vitro* and functionally complements SpoT in *E. coli* ^4^ Furthermore, *Meshl* deletion in *Drosophila* displays impaired starvation resistance, a susceptibility that is fully rescued by *Meshl* expression^4^. Despite these supporting lines of evidence, the existence of a ppGpp-mediated stringent response pathway in metazoans has serious impediments, as neither ppGpp nor its synthetase has been discovered in metazoans, and no physiological substrate for MESH1 has been reported.

## MESH1 is an efficient NADPH phosphatase

We reasoned that MESH1 may function through alternate metabolic substrate(s) from ppGpp in mammalian cells. Consequently, we tested purified-recombinant human MESH1 (hMESH1) against a set of common metabolites, such as UDP-glucose (uridine diphosphate glucose), GDP (guanosine diphosphate), CDP (cytidine diphosphate), pyrophosphate, creatine phosphate, GDP-fucose, thiamine pyrophosphate and various form of inositol phosphates, but failed to detect any activity (data not shown). We then examined metabolites with a similar molecular architecture to ppGpp. The metabolite NADPH shares many similarities with ppGpp including a purine nucleoside, a 2’ phosphate group, and a 5’ pyrophosphate group (Fig. 1a). Although NADPH differs from ppGpp in that it contains a 2’ phosphate instead of a 3’ pyrophosphate in ppGpp, the crystal structure of the bifunctional RelA/SpoT homologue from *Streptococcus dysgalactiae subsp. equisimilis* captured an unusual ppGpp derivative, GDP-2’,3’ - cyclic monophosphate, in the active site of the hydrolase domain ^5^, suggesting that the enzyme may accommodate a 2’-substituted phosphate group ^6^. Based on the observation that SpoT catalyzes the hydrolysis of the 3’-pyrophosphate group of ppGpp, we predicted that MESH1 would similarly hydrolyze the 2’-phosphate group of NADPH to yield NADH and an inorganic phosphate (Fig. 1a).

**Figure. 1.**
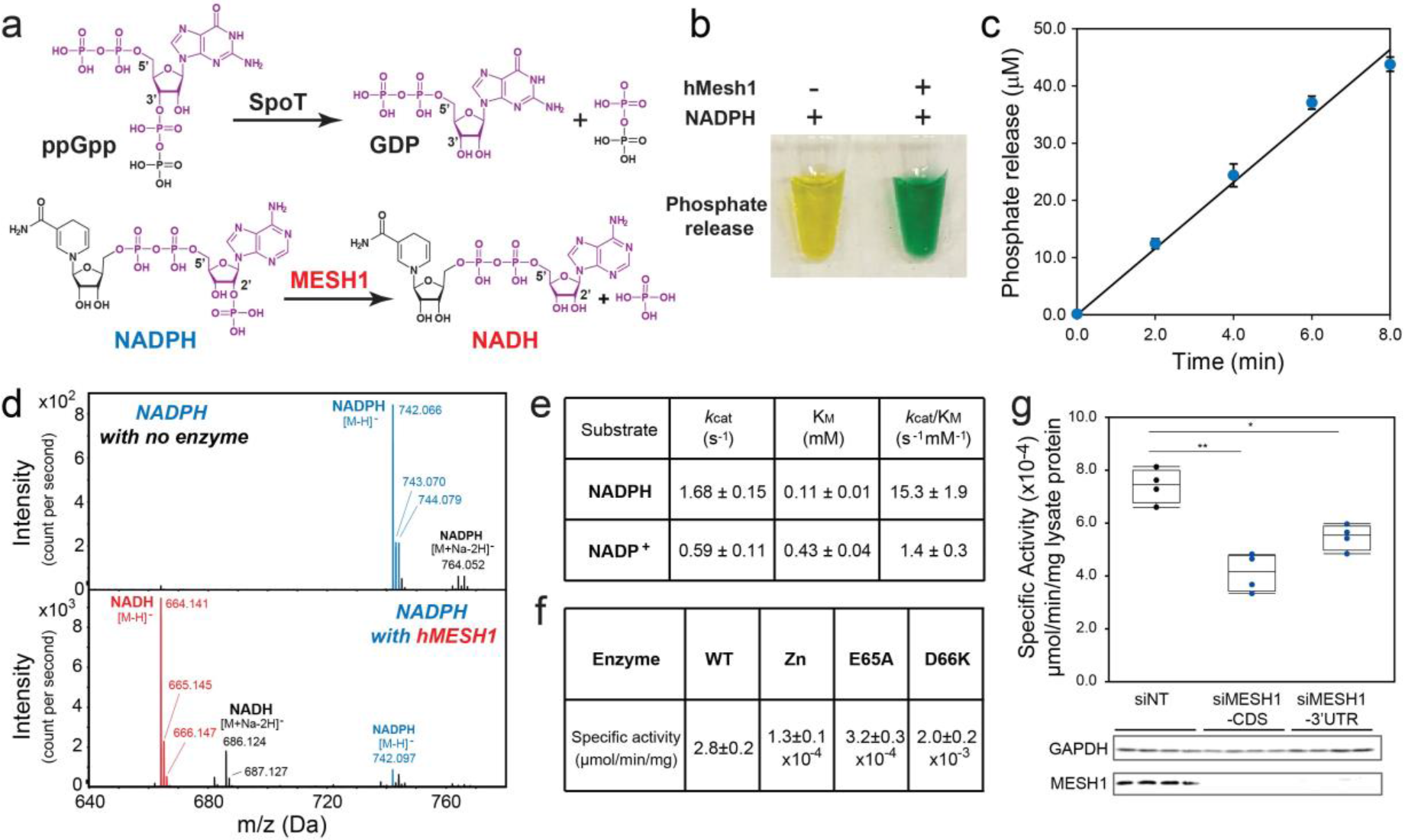
MESH1 is a mammalian NADPH phosphatase. **a**, Chemical similarity between NADPH and ppGpp and proposed chemical reaction of MESH1. **b**, Detection of the hMESH1-catalyzed phosphate release from NADPH by the colorimetric malachite green assay. The malachite green reagent changes from yellow to green in the presence of inorganic phosphate. **c**, Linear product accumulation of the NADPH dephosphorylation reaction catalyzed by hMESH1. **d**, Validation of the formation of NADH by mass spectrometry analysis. **e**, Enzymatic characterization of hMESH1 toward NADPH and NADP^+^. **f**, Effects of Zn^2+^ substitution and active site mutation on the specific activity of purified recombinant hMESH1. **g**. hMESH1 is a significant contributor to the NADPH phosphatase activity in RCC4 lysates. Representative samples were analyzed by Western blot to confirm knock-down efficiency. *n*=4 for each condition. One-way ANOVA with Tukey HSD post-hoc test. \**P*<0.05, \*\**P*<0.01

Indeed, treatment of NADPH with hMESH1 readily released inorganic phosphate, yielding a green solution in the malachite green assay ^7^ (Fig. 1b). Importantly, the phosphate accumulation was linear over time (Fig. 1c), reflecting continuous enzymatic turnover of NADPH by hMESH1. Furthermore, mass spectrometry analysis revealed a peak at m/z 664.141 as [M-H]^−^, verifying the product as NADH (Fig. 1d). We then analyzed the steady state kinetics parameters of hMESH1 toward NADPH (Fig. 1e). hMESH1 is an efficient NADPH phosphatase, with a catalytic efficiency (k_cat_/*K*_M_) of 15.3 ± 1.9 s^−1^mM^−1^; its *K*_M_ value of 0.11 ± 0.01 mM is on par with that from other reported cellular enzymes that utilize NADPH (e.g., *K*_M_ value of 0.11 mM for the human phagocytic NADPH oxidase ^8^), supporting its role as a physiologically relevant NADPH phosphatase in cells. While hMESH1 also displays measurable activity towards NADP^+^ *in vitro*, it is ~10-fold less efficient (k_cat_/*K*_M_ = 1.4 ± 0.3 s^−1^mM^−1^), and its *K*_M_ value of 0.43 ± 0.03 mM is significantly higher than the estimated cytosolic concentration of NADP^+^, rendering hMESH1 an ineffective phosphatase for NADP^+^ in cells (Fig. 1e).

Previous biochemical and structural analysis has revealed MESH1 as a Mn^2+^-dependent enzyme^4^. Accordingly, we found that the NADPH phosphatase activity of hMESH1 is compromised when Mn^2+^ was substituted with other metal ions, such as Zn^2+^ (Fig. 1f). Likewise,
mutations of hMESH1 residues near the Mn^2+^ ion in the active site, including E65A and D66K, also severely compromised the NADPH phosphatase activity in enzymatic assays (Fig. 1f).

In order to determine whether MESH1 is a significant contributor to the cellular NADPH phosphatase activity, we measured the enzymatic activity in cell lysates extracted from two human cell lines, RCC4 cells (a human clear cell renal carcinoma cell line) and HEK-293T cells. We found that the *MESH1*-silenced cell lysates had a substantial decrease of NADPH phosphatase activity (Fig. 1g, Extended Data Fig. 1a). Conversely, overexpression of wild-type hMESH1, but not the catalytically inactive E65A hMESH1 mutant, significantly enhanced the cellular NADPH phosphatase activity (Extended Data Fig. 1b). Taken together, these observations verify hMESH1 as a significant contributor to the NADPH phosphatase activity in human cells. It is worth noting that the siRNA of *MESH1* did not suppress NADPH phosphatase activity completely, indicating the possible existence of another enzyme that also contributes to NADPH phosphatase activity in mammalian cells. One possible candidate is the NADP(H) phosphatase activity previously observed in rat liver lysates ^9^, though the reported enzymatic activity favors NADP^+^ and the identity of the enzyme has remained unidentified in animal cells^10^. Therefore, to the best of our knowledge, the NADPH phosphatase activity of MESH1 is distinct from the previously observed activity and represents the first description of an NADPH phosphatase in human cells.

## Molecular recognition of NADPH by hMESH1

In order to visualize the molecular details of the NADPH-recognition, we determined the co-crystal structure of the hMESH1-NADPH complex by using the catalytically compromised D66K mutant enzyme and by substituting the catalytic Mn^2+^ ion with Zn^2+^. The structure was refined to 2.1 Å resolution (**Extended Data Table 1**). The crystallographic asymmetric unit consists of two protomers of hMESH1 (Extended Data Fig. 2), each adopting a compact fold of ten α-helices and α short β-hairpin (Fig. 2a) previously observed in the structure of apo hMESH1 ^4^ While the active sites of both protomers were occupied by the adenosine portion of the substrate, the molecular recognition of the entire NADPH molecule, including that of the nicotinamide moiety, was only visible in one of the two protomers, which is described below.

**Figure. 2.**
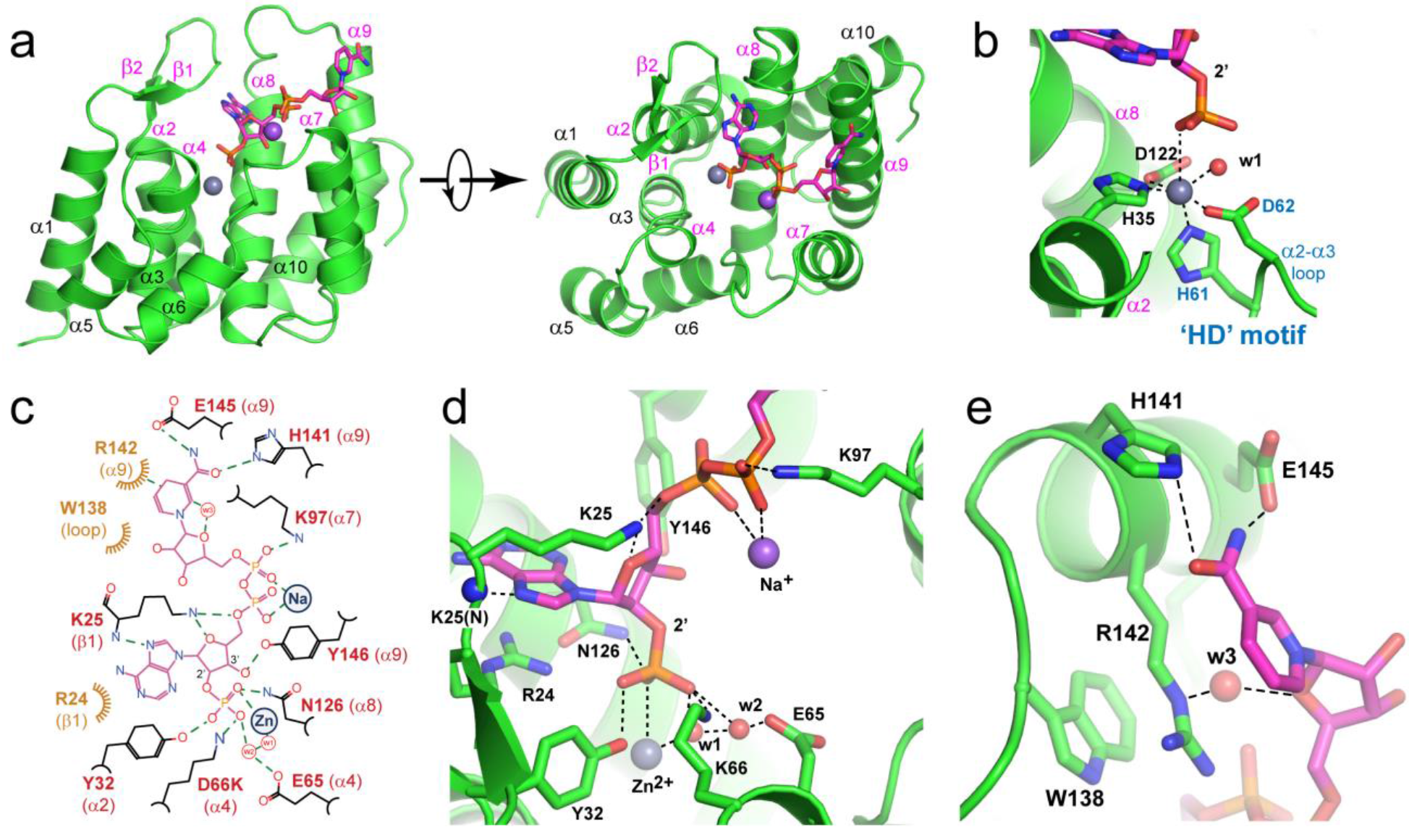
Structure of the hMESH1 (D66K)-NADPH complex. **a**, The architecture of the hMESH1 active site. MESH1 is shown in the ribbon diagram, NADPH in the stick model, and the Zn^2+^ (grey) and Na^2+^ (pink) ions are shown as spheres. Secondary structures are labeled, with MESH1-NADPH-interacting motifs annotated in magenta. **b**, Coordination of the active site Zn^2+^ ion in a distorted octahedral geometry. NADPH and sidechains of the Zn^2+^-binding residues are shown in the stick model. The zinc ion (grey) and its coordinating water molecule (red) are shown as spheres. The signature ‘HD’ motif is annotated in blue. **c**, A schematic illustration of the NADPH recognition by MESH1. Polar interactions are denoted with dashed lines, and vdW contacts are shown with curved lines. The location of the metal ions and NADPH-interacting water molecules are denoted as circles. **d**, Molecular recognition of the 2’-phospho-adenosine diphosphate moiety. **e**, Molecular recognition of the nicotinamide riboside moiety.

The catalytic site of hMESH1 is formed at the center of the helical architecture, surrounded by the short β-hairpin (β1 and β2), α2, the α3-α4 loop, α4, α7, α8, the α8-α9 loop, and α9 (Fig. 2a). Anomalous scattering revealed the presence of a single Zn^2+^ ion in the active site, which substitutes the catalytic Mn^2+^ ion reported in the apo hMESH1 structure (Extended Data Fig. 2b). The Zn^2+^ ion is hexa-coordinated in a distorted octahedral geometry by H35 of α2, H61 and D62—the signature HD motif of the α2-α3 loop that defines this family of enzymes, D122 of α8, the catalytic water molecule, and the 2’-phosphate group of NADPH (Fig. 2b).

In addition to coordinating the active site metal ion, the 2’-phosphate group of NADPH forms additional polar interactions with Y32 of α2, K66 of α4, and N126 of α8 that are located one layer above the equatorial plane of the zinc ligands (Fig. 2c,d). K66, the mutated residue substituting D66 in the WT enzyme, not only significantly diminishes the catalytic activity, but also forms a direct salt bridge with the 2’-phosphate group of NADPH, a likely contributor to the successful capture of NADPH in the co-crystal structure. Another catalytically important residue, E65 of α4, the neighboring residue of D66, is located over 4 Å away from the catalytic water molecule, and its indirect interactions with the catalytic water molecule and the 2’phosphate of NADPH are bridged by a second water molecule in the active site (Fig. 2c, d).

The NADPH ribose group and its 5’ pyrophosphate group are extensively recognized (Fig. 2c,d). The 3’-hydroxyl group and the 4’-oxygen atom form hydrogen bonds with Y146 of α9 and K25 of β1, respectively. The 5’ diphosphate group adopts a tight turn, aided by salt bridges with K97 of α7, K25 of β1 and a sodium ion. While sodium is an unusual cation to coordinate the NADPH diphosphate, our purification procedure and crystallization condition lack divalent cations other than Zn^2+^ but contain over 600 mM sodium chloride. Given this density lacks anomalous Zn^2+^ signals (Extended Data Fig. 2b), it is interpreted as a sodium ion, though Mg^2+^ or Ca^2+^ might be more suitable ions *in vivo*. The adenine moiety of NADPH is largely coordinated by π-stacking with R24 of β1 and a direct hydrogen bond between its N7 atom and the amide group of K25 of the β1 loop (Fig. 2c,d). As both of these interactions are also found in a guanine base, hMESH1 lacks the ability to differentiate the adenine nucleotide from the guanine nucleotide as was found in ppGpp.

The ribose ring of the nicotinamide riboside is indirectly recognized by a water mediated hydrogen bond of its ring oxygen atom with R142 of α9, whereas the nicotinamide moiety is supported by a pi-stacking network with W138 and R142 that emanate from the α8-α9 loop and from α9 (Fig. 2c,e). The terminal amide group is additionally buttressed by two hydrogen bonds with H141 and E145 of α9. These structural observations reinforce the notion that hMESH1 is a *bona fide* NADPH phosphatase.

## MESH1 regulates cellular NADPH levels and ferroptosis

After revealing the molecular basis of the NADPH recognition by hMESH1 and establishing hMESH1 as a significant contributor to the cellular NADPH phosphatase activity, we examined the functional impact of manipulating MESH1 in human cells on NADP(H) and ferroptosis. Ferroptosis is an iron-dependent death under oxidative stresses^11^ which can be mitigated by a higher concentration of NADPH^12^. As expected, overexpression of wild type hMESH1 (MESH1-WT), but not an enzymatically deficient mutant (MESH1-E65A), significantly lowered intracellular NADP(H) (Extended Data Fig. 3a, b). However, the reduction of *MESH1* expression by RNAi does not statistically alter the NADP(H) steady state level in cells (Extended Data Fig 3c). We then asked whether MESH1 depletion could regulate NADP(H) levels and cell viability when treated with erastin, which is known to diminish intracellular NADP(H)^12,13^ and trigger ferroptosis^11^. We examined the NADPH concentrations of *MESHl*-silenced human cells before and after treatment with erastin. While erastin treatment dramatically reduced the NADPH level in the control (siNT) cells, the NADPH level was sustained at a significantly higher level in *MESH1*-silenced (siMESH1) cells (Fig. 3a). Accordingly, the RNAi-mediated *MESH1*-depletion significantly increased cell survival in the presence of erastin over control cells (Fig. 3b). Such an effect appeared to be general, as *MESH1* depletion similarly rescued cell death in multiple cell lines under erastin (Extended Data Fig. 3d-f). Importantly, the resistance to erastin-induced ferroptosis was eliminated by overexpression of WT *MESH1,* but not the catalytically compressed E65A mutant of *MESH1* (Fig. 3b), indicating that the enhanced cell viability is directly associated with the catalytic activity of hMESH1.

**Figure. 3.**
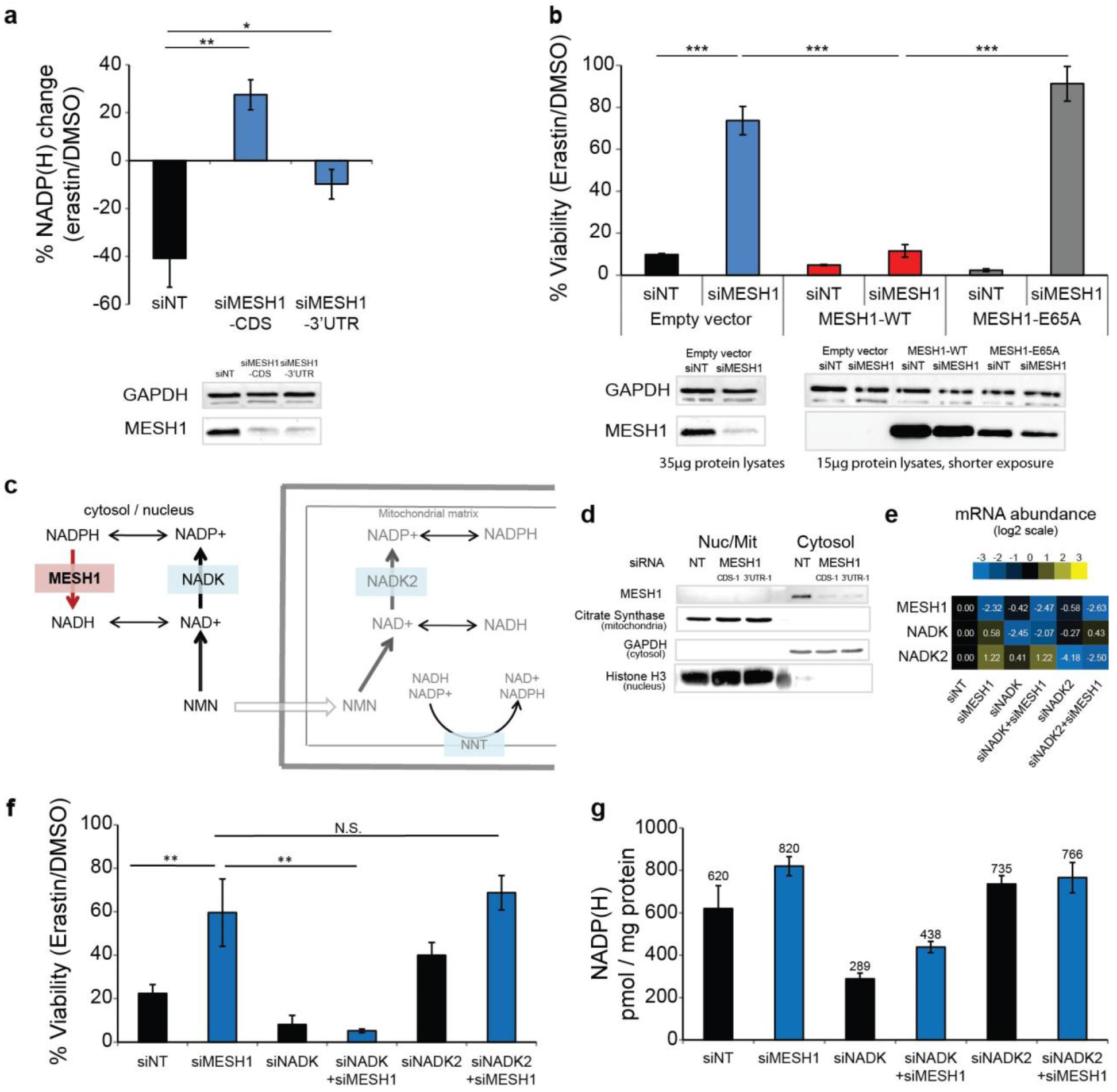
MESH1 regulates cellular NADP(H) levels and ferroptosis. **a**, The percentage change of NADP(H) after one day of erastin treatment in RCC4 cells with non-targeting (NT) siRNA or 2 distinct MESH1-targeting siRNAs. MESH1 knock-down efficiency was validated by Western blot. **b**, Percentage of cell viability of RCC4 cells after erastin treatment for one day. Lentivirus with empty vector, MESH1-WT or MESH1-E65A, were stably induced into RCC4 prior to siRNA transfection. The knockdown and overexpression efficiency was validated by Western blot. If not specified, siMESH1 indicates siMESH1-CDS. **c**, A schematic model of NADH/NADPH metabolism elucidating the proposed role of MESH1 as a cytosolic NADPH phosphatase. **d**, Cell fractionation separating cytosol from nucleus and mitochondria fraction, showing that MESH1 is specifically enriched in cytosolic compartment. **e**, rt-qPCR to quantify the normalized levels of MESH1, NADK and NADK2 in cells with indicated siRNAs. **f**, NADK knockdown eliminates the survival advantage seen in MESH1 silenced cells upon erastin treatment, bars 2 and 4, while mitochrondrially localized NADK2, bars 5 and 6, does not. **g**, Co-silencing of both siNADK and MESH1 reduces measured NADP(H) levels. (**f-g**) RCC4 cells after siRNA transfection for 2 days and erastin incubation for one day. *n*=3-5 per condition. Oneway ANOVA with Tukey HSD post-hoc test **(a)**, Two-way ANOVA with Tukey HSD post-hoc test (b, f). Error bar indicates s.d.m. \**P*<0.05; \*\**P*<0.01; \*\*\**P*<0.005; N.S., not significant;

Because NADPH is a central metabolite, directly assessing the effects of MESH1 on downstream pathways proved difficult. Thus, to further establish that the enhanced cell survival under oxidative stress is due to a higher sustained level of NADPH, but not due to accumulation of an unidentified substrate of hMESH1, we also tested whether the survival advantage of MESH1 depletion is reversed by simultaneous depletion of NAD(H) kinases that convert NAD(H) to NADP(H). Human cells have two NAD kinases, NADK and NADK2, that are predominantly located in cytosol or mitochondria^14,15^, respectively (Fig. 3c). As the cytosolic and mitochondrial pools of NADP(H) and NAD(H) are compartmentalized in mammalian cells due to the impermeability of these molecules across the mitochondrial membranes^16^, we reasoned that the removal of the NAD kinase either from the cytosol (NADK) or mitochondria (NADK2) would reduce the distinct pools of NADP(H) and compromise the ferroptosis survival phenotypes of *MESH1*-silenced cells. Indeed, when the gene encoding the cytosolic enzyme NADK was silenced, the survival benefit of silencing *MESH1* was largely eliminated (Fig. 3e-f, Extended Data Fig. 3g) and the intracellular NADP(H) is largely depleted (Fig. 3g). In contrast, silencing the gene encoding the mitochondrial enzyme NADK2 did not affect the MESH1-mediated ferroptosis survival or NADP(H) (Fig. 3e-g, Extended Data Fig. 3g)), suggesting its limited role in this process. Consistent with these observations, we found that MESH1 was predominately enriched in the cytosolic, but not the mitochondrial or nuclear pools of proteins, by fractioning the individual cellular components^17^ (Fig. 3d). Taken together, our results establish MESH1 as a cytosolic NADPH phosphatase, and depletion of its activity promotes stress survival for mammalian cells under ferroptosis-inducing conditions by sustaining a higher level of NADPH.

## *MESHl*-silencing induces an extensive transcriptional response

After demonstrating that *MESH1* silencing promotes mammalian cell survival under oxidative stress, we examined the transcriptional profiles of *MESH1*-silenced cells to determine if the signatures of this stress survival phenotype are similarly observed in other mammalian stress survival pathways, such as the integrated stress response (ISR) ^18^.

Our transcriptome analysis using two independent siRNAs directed to *MESH1* revealed that *MESH1*-silencing triggered extensive transcriptional responses in RCC4 cells. Further Gene Ontology (GO) analysis of differentially expressed genes showed that *MESH1*-silencing repressed pathways relevant to cell cycle progression and DNA replication (Fig. 4a), with a striking similarity to those observed in the bacterial stringent response^19,20^ and in *Mesh1* deficient Drosophila^4^. The inhibition of cell cycle regulated genes, *CDK2* (Cyclin Dependent Kinase 2), *RRM2* (Ribonucleotide Reductase Regulatory Subunit M2), and *E2F1* (E2F transcription factor 1) were validated via rt-qPCR (Fig. 4b-d).

**Figure. 4.**
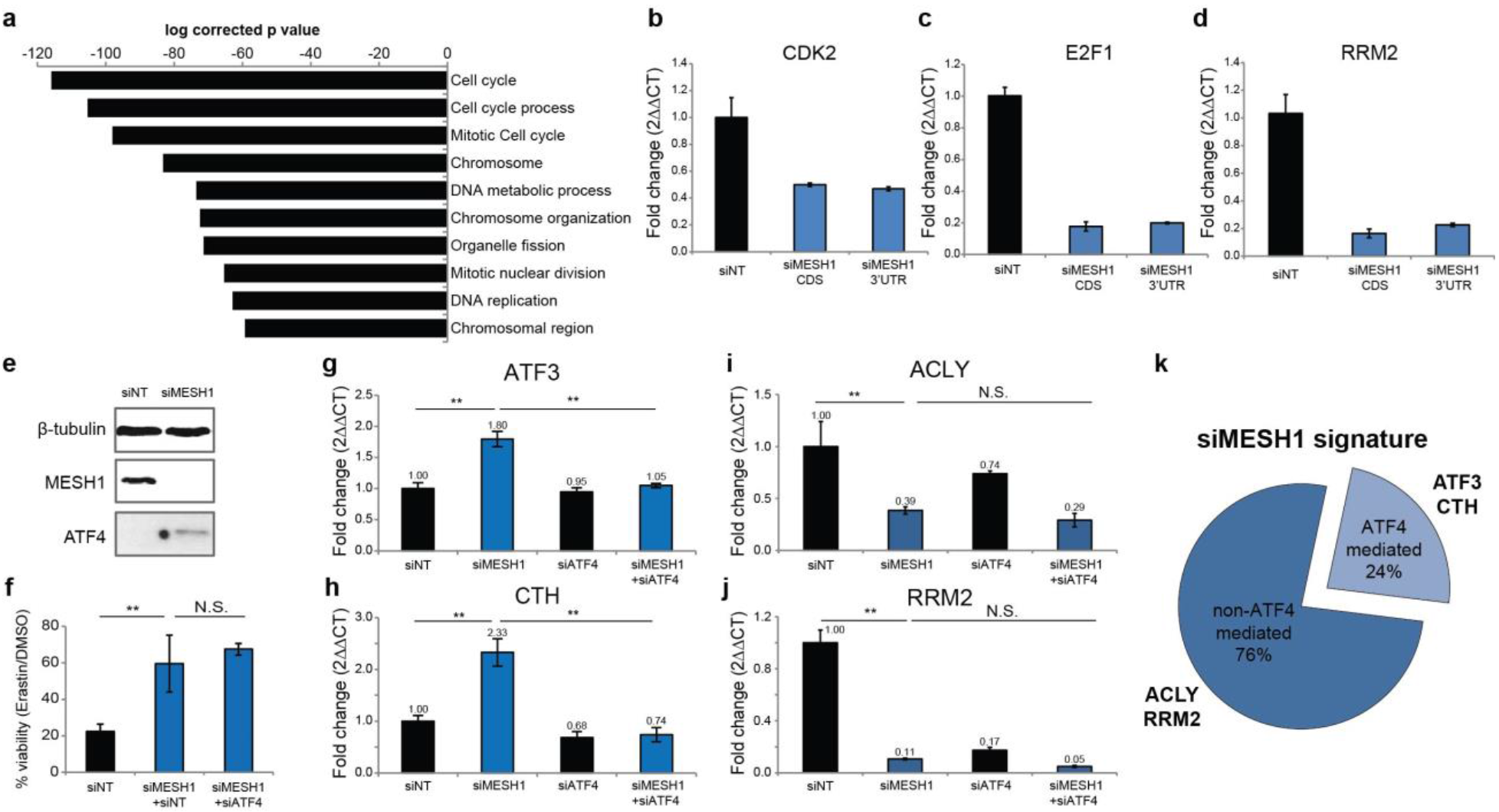
The transcriptional reprogramming upon *MESH1* removal. **a**, Top 10 commonly inhibited Gene Ontology processes of the transcriptome of *MESH1*-silenced RCC4 cells. *MESH1* is silenced by siMESH1-CDS or siMESH1-3’UTR and profiled by RNA-seq. RNA was collected after siRNA transfection for 3 days. **b-d**, rt-qPCR validation of mRNA abundance of *CDK2, E2F1* and *RRM2.* e, ATF4 is upregulated in *MESH1*-silenced RCC4 cells. f, *ATF4-* silencing did not influence resistance to erastin induced by *MESH1*-silencing. **g-j**, mRNA level of ATF4-mediated genes (**g, h**) and non-ATF4-mediated genes (**i, j**). mRNA abundance is normalized by β-actin and presented in fold change (2^▵▵CT^) (**b-d**, **g-j**). **k**. The majority of the transcriptional signature of MESH1 removal was not mediated by ATF4, estimated by transcriptional profiling of *MESH1*-silenced, *ATF4*-silenced, *MESH1-ATF4*-silenced, and control (siNT) RCC4 cells using cDNA microarrays. Two-way ANOVA with Tukey-HSD post-hoc, *n*=3 (**g-j**). Error bar indicates s.d.m. for triplicates (**b-d**, **g-j**). \**P*\*0.05, \*\**P*<0.01, \*\*\**P*<0.005; N.S., not significant. siMESH1 signature is defined as differential gene expression with fold change >1.74 with t-test *P*<0.05. ATF4-mediated genes are defined as having a siMESH1 signature that are reversed upon additional *ATF4-silencing* with fold change > 1.41 and t-test *P*<0.05 compared to *MESH1*-silencing toward the direction of siNT samples (**k**).

Interestingly, MESH1-silencing induced the expression of ATF4 (Activating transcription factor 4, also known as cAMP-responsive element binding protein 2) (Fig. 4e)—a key regulator of the integrated stress response—and its target genes, such as ATF3 (Activating transcription factor 3) and CTH (Cystathionine gamma-lyase)^21^ (Fig. 4g, h), suggesting that MESH1-silencing also triggered a known mammalian stress response. In order to understand the relationship between the *MESH1*-silencing-induced stress response and the integrated stress response, we simultaneously silenced *MESH1* and *ATF4* and compared the resulting cellular phenotype and transcriptional changes with those of the *MESH1*-silenced cells. We found that *ATF4*-silencing did not alter the stress survival phenotype of *MESH1*-silenced cells under ferroptosis-inducing conditions (Fig. 4f). In addition, while the *ATF4*-silencing abolished the induction of known *ATF4* target genes, including *ATF3* and *CTH* (Fig. 4g, h), it did not affect the transcriptional profile of other *MESH1*-silenceing responsive genes such as *ACLY* (ATP Citrate Lyase) and *RRM2* (Fig. 4i, j). In general, only 24% (124 out of 525 genes) of the *MESH1*-silencing signature was affected by simultaneous ATF4 silencing (Fig. 4k). These data argue that although the *MESH1*-silencing-induced stress response may function in part through genes in the integrated stress survival pathway, it has a distinct transcriptional profile and regulatory mechanism and functions independently of the integrated stress survival pathway to promote stress survival under ferroptosis-inducing conditions.

In summary, we found that MESH1, the metazoan homolog of SpoT, is a cytosolic NADPH phosphatase in human cells. *MESH1*-silencing preserves NADPH, which contributes to cell survival under ferroptosis-inducing conditions. Although the transcriptional profile of *MESH1*-silenced cells partially overlaps with that of the integrated stress response, the stress survival phenotype of *MESH1*-silencing is independent of ATF4, a key mediator of the integrated stress response pathway. Given the striking similarity of transcriptional changes caused by *MESH1*-silencing with that of the bacterial stringent response, we suggest that >MESH1 plays a key role in a novel mammalian stress-response pathway that resembles the bacterial stringent response.

## Methods

### Purification of recombinant hMESH1

The gene encoding hMESH1 was codon optimized for *E. coli* expression, synthesized, and cloned into a modified pET28a vector as a C-terminal fusion to the His_6_-tagged SUMO protein. Cultures of transformed *E. coli* strain BL21(DE3)* were grown to an optical density at 600 nm (OD_600_) between 0.4 and 0.5 and induced with 1 mM Isopropyl β-D-1- thiogalactopyranoside (IPTG) at 37 °C for 2 hours. Following cell lysis, the target protein was purified using Ni-NTA affinity chromatography following standard protocols (Qiagen). The SUMO tag was cleaved using the SENP1 protease, and both the tag and protease were removed by a second round of Ni-NTA chromatography. The target protein was further purified using size-exclusion chromatography (Superdex 75; GE life sciences) in a buffer containing 50 mM Tris pH 8.0, 200 mM NaCl, 0.1% 2-mercaptoethanol. Mutants of hMESH1 were generated using the QuikChange site directed mutagenesis kit (Agilent) and prepared using the same procedure.

### Mass spectrometry

RPLC-ESI/MS/MS was performed using a Shimadzu LC system (comprising a solvent degasser, two LC-10A pumps and a SCL-10A system controller) coupled to a high-resolution TripleTOF5600 mass spectrometer (Sciex). LC was operated at a flow rate of 200 μl/min with a linear gradient as follows: 100% of mobile phase A was held isocratically for 2 min and then linearly increased to 100% mobile phase B over 5 min and held at 100% B for 2 min. Mobile phase A was a mixture of water/acetonitrile (98/2, v/v) containing 0.1% acetic acid. Mobile phase B was a mixture of water/acetonitrile (10/90, v/v) containing 0.1% acetic acid. A Zorbax SB-C8 reversed-phase column (5 μm, 2.1 × 50 mm) was obtained from Agilent. The LC eluent was introduced into the ESI source of the mass spectrometer. Instrument settings for negative ion ESI/MS and MS/MS analysis of lipid species were as follows: Ion spray voltage (IS) = −4500 V; Curtain gas (CUR) = 20 psi; Ion source gas 1 (GS1) = 20 psi; De-clustering potential (DP) = −55 V; Focusing potential (FP) = −150 V. Data acquisition and analysis were performed using the Analyst TF1.5 software (Sciex).

### Enzymology

Enzymatic assays for hMESH1 were performed in a buffer containing 50 mM Tris pH 8, 200 mM NaCl, and 1 mM MnCl_2_ or ZnCl_2_. Assays for NADPH and NADP^+^ were carried out using 50 nM or 500 nM enzyme, respectively. Reactions were stopped by the addition of formic acid (3 M final concentration). The amount of released phosphate was assessed using the malachite green reagent as previously described ^7^. Values for K_M_ and V_max_ were calculated from the Michaelis-Menten equation. Specific activities of purified hMESH1 mutants or cell lysates were determined using 1 mM substrate, with the reaction time varying from 15-90 minutes.

### X-ray crystallography

Crystals of hMESH1 bound to NADPH were formed using the hanging drop vapor diffusion method. Immediately prior to crystallization, hMESH1 was treated with 10 mM EDTA for 30 min and exchanged into a buffer containing no metal before addition of 1 mM ZnCl_2_. Diffracting quality crystals were obtained by mixing equal volumes of the protein solution (9 mg/mL hMESH1, 100 mM NADPH, 50 mM Tris pH 8.0, 200 mM NaCl and 0.1% 2-mercaptoethanol) with the mother liquor solution (200 mM ammonium acetate, 100 mM sodium citrate, and 25% PEG 4000) by streak-seeding with apo hMesh1 crystals. To improve NADPH occupancy, crystals were additionally soaked in the mother liquor solution containing 750mM NADPH for 1 hour prior to cryoprotection in a solution containing 525 mM NADPH, 10 mM ZnCl_2_, 20% ethylene glycol, and 70% precipitant solution and flash frozen in liquid nitrogen.

X-ray data were collected at Southeast Regional Collaborative Access Team (SER-CAT) 22-BM beamline at the Advanced Photon Source, Argonne National Laboratory, processed using XDS ^22^, and scaled using the UCLA diffraction anisotropy server ^23^. The structure was determined by molecular replacement with the program Phaser ^24^ and by using the previously reported structure of apo hMESH1 (3NR1) as the search model. Iterative model building and refinement was carried out using COOT ^25^ and PHENIX ^26^.

### *MESH1*-silencing using RNAi

Non-targeting siRNA (siNT) was purchased from Qiagen (AllStars Negative Control siRNA, SI03650318). Other siRNA includes: siMESH1-CDS (target sequence GGGAAUCACUGACAUUGUG, D-031786-01, Dharmacon), siMESH1-3’UTR (target sequence CTGAAGGTCTCCTGCTAACTA, SI04167002, Qiagen), siATF4 (target sequence GAUCAUUCCUUUAGUUUAG, CAUGAUCCCUCAGUGCAUA, GUUUAGAGCUGGGCAGUGA, CUAGGUACCGCCAGAAGAA, M-005125-02, Dharmacon), siNADK (target sequence UGAAUGAGGUGGUGAUUGA, CGCCAGCGAUGAAAGCUUU, GAAGACGGCGUGCACAAU, CCAAUCAGAUAGACUUCAU, M-006318-01, Dharmacon), and siNADK2 (target sequence GCAAUU GCUU C GAU GAUGA, GAGAAUUGGUAGAGAAAGU, UGUGAAAGCUGGAC GGAU A,UUUGAUUACUGGCGAGAAU, M-006319-01, Dharmacon). If not specified, siMESH1 indicates siMESH1-CDS. The efficacies of these siRNA were accessed by rt-qPCR and/or Western blots. For enzymology, 8 × 10^5^ RCC4 cells were seeded in a 100mm plate, and transfected after one day of growth with 600 pmole of siRNA and 40 μL Lipofectamine RNAiMAX (ThermoFisher Scientific, #13778150) for 72 hours before the collection of cell lysates. For NADP(H) measurement, 8 × 10^4^ cells were seeded per well of 6-well plate with 40 pmole of siRNA and transfected using 3μL of Lipofectamine RNAiMAX for 48 hours before erastin treatment for one day. For viability assays, 2,800 RCC4 cells were seeded per well on 96-well plates with 5 pmole of siRNA and 0.4 μL Lipofectamine RNAiMAX for at least 48 hours before erastin treatmet for one day. To collect RNA or protein, 10^5^ RCC4 cells were seeded in a well of a 6-well plate with 40 pmole of siRNA and 3μL of Lipofectamine RNAiMAX for 72 hours before collection.

### MESH1-overexpression

Stable RCC4 cell lines with MESH1-WT or MESH1-E65A expression were generated using lentiviral vector pLX302^27^ (gift from David Root, Addgene plasmid #25896) without expression of the V5 tag. Briefly, virus was generated by transfecting HEK-293T cells with a 0.1:1:1 ratio of pMD2.G: psPAX2: pLX302 with TransIT-LT1 Transfection Reagent (Mirus, MIR2305). pMD2.G and psPAX2 were gifts from Didier Trono (Addgene plasmid #12259 and #12260, respectively). Virus was collected 48 hours after transfection. Stable cell lines were generated by adding 200 μL virus to a 60 mm dish of RCC4 cells with 8 μg/mL polybrene. 1 μg/mL puromycin was used for selection. Complete death in blank infection dishes was used to determine the success of infection and the length of puromycin selection. The efficiency of overexpression was determined by Western blotting.

Transient overexpression of MESH1 in HEK-293T cells for NADP(H) content assay or enzymology assay was achieved by using the pCMV6-neo vector (OriGene, SC334209). Briefly, 2 × 10^5^ HEK-293T cells were seeded in one well of a 6-well plate for 24 hours, and then 1 pg of plasmid (pCMV6-neo empty vector or with MESH1-WT or MESH1-E65A) was transfected with TransIT-LT1 Transfection Reagent (Mirus) for additional 48 hours before collection.

### Cell culture

RCC4 cell line was provided by Denise Chan (University of California, San Francisco, San Francisco, CA), and was authenticated by DDC (DNA Diagnostics Center) Medical using the short tandem repeat method in November 2015. HEK-293T, H1975, and MDA-MB-231 were obtained from the Duke Cell Culture Facility. All cells were cultured in DMEM with 4.5 g/L glucose and 4 mM Glutamine (11995-DMEM, ThermoFisher Scientific) and 10% heat-inactivated fetal bovine serum (Hyclone # SH30070.03HI) in humidified incubator, 37°C with 5% CO2.

### NADP(H) measurement

Amplite colorimetric NADPH assay kit (#15272, ATT Bioquest) was used to measure NADP(H) (both NADP^+^ and NADPH) in this study. Briefly, cells were seeded into a 6-well plate with siRNA or plasmid as described. NADP(H) was collected using 100 μL of the provided lysis buffer. NADP(H) content was measured and normalized by protein content of the lysate, quantified by BCA assay. In cells treated with erastin, the NADP(H) change is normalized with NADP(H) measured in DMSO-treated cells.

### Protein lysate collection and Western blots

Cells lysates were collected in buffer with Tris 50mM pH8.0 and NaCl 200mM (for enzymology), NADP(H) lysis buffer (for NADP(H) measurement), or Radioimmunoprecipitation assay (RIPA) buffer (Sigma, R0278) with protease inhibitors (Roche, 11836170001). For Western blots, 15-50 αg of lysates were loaded on SDS-PAGE gels, semi-dry transferred to PDVF membrane, blocked with 5% milk in TBST with 0.1% Tween-20, then incubated with primary antibodies overnight at 4°C. Anti-GAPDH antibody (Santa Cruz, sc-25778), anti-MESH1 antibody (Abcam, ab118325), anti-β-tubulin antibody (Cell Signaling Technology, #2128), anti-citrate synthetase antibody (GeneTex, GTX110624), anti-histone-H3 antibody (Cell Signaling Technology, #4499), anti-ATF4 antibody (Santa Cruz, sc-200), anti-mouse-IgG HRP (Cell Signaling Technology, #7076), and anti-rabbit-IgG HRP (Cell Signaling Technology, #7074) were used in this study. The images were developed by SuperSignal West Pico PLUS Chemiluminescent substrate (ThermoFisher, #34577) or Amersham ECL Prime Western Blotting Detection Reagent (GE Healthcare Life Sciences, RPN2232) and exposed in ChemiDoc imaging system (Biorad).

### Quantitative real-time PCR

RNA was extracted using the RNeasy kit (Qiagen, 74104) following manufacturer’s instructions. 500 ng of total RNA with or without reverse transcriptase were prepared using a SensiFast cDNA synthesis kit (Bioline, BIO-65054) for real-time PCR comparison with Power SYBRGreen Mix (ThermoFisher Scientific, 4367659). Primers were designed across exons whenever possible using PrimerBot! developed by Jeff Jasper at Duke University. The product of PCR was validated for specificity by DNA electrophoresis.

### Transcriptome analysis

Total RNA was isolated as detailed above. RNA quality was assessed using an Agilent BioAnalyzer (Agilent). For RNAseq, 1 μg of total RNA was used to generate a cDNA library with Illumina TruSeq Stranded mRNA LT Sample Prep Kit – Set A (Illumina, RS-122-2101) according to the manufacturer’s instructions. The library was pooled and sequenced using Illumina HiSeq 4000 with single-end 50 bp read length at The Sequencing and Genomic Technologies Shared Resource of Duke Cancer Institute. The data was analyzed using TopHat^28^ and HTSeq ^29^ with USCS hg19 as the reference genome. The differential analysis was performed using DESeq2 ^30^. The genes that are differentially expressed (adjusted p value <0.05) in both siMESH1-CDS and siMESH1-3’UTR were selected and analyzed using MSigDB (Molecular Signatures Database) database v6.1 and web site v6.3 with C5 (GO gene ^31,32^. for Gene Ontology analysis. For cDNA microarray, cDNA was amplified from 200 ng RNA with Ambion MessageAmp Premier RNA Amplification kit (ThermoFisher Scientific, AM1792). The gene expression signatures were interrogated with Affymetrix U133A genechips and normalized by the RMA (Robust Multi-Array) algorithm. cDNA synthesis and microarray interrogation was performed by the Duke Microarray Core. The influence of silencing MESH1 and ATF4 on gene expression was derived by the zero transformation process, in which we compared the gene expression level with siMESH1 or siMESH1+siATF4 to the average of gene expression level in control (siNT) samples. Significant differentially-expressed genes (t test, *P*<0.05) were identified and a selection of genes were validated by quantitative rt-PCR.

## Acknowledgements

This work was supported in part by DOD grants (W81XWH-17-1-0143, W81XWH-15-1-0486 to J.T.C.), NIH grant GM115355 (to P,Z.), the Duke Bridge Fund, Duke Cancer Institute (DCI) Discovery fund and DCI pilot fund. X-ray diffraction data were collected at the Southeast Regional Collaborative Access Team (SER-CAT) 22-ID beamline at the Advanced Photon Source, Argonne National Laboratory, supported by the US Department of Energy, Office of Science and the Office of Basic Energy Sciences under Contract number W-31-109-Eng-38. We appreciate the great suggestions and help from W.H. Yang, J. Zhao, and the X. Shen lab and J. Zhao

## Author Information

### Contributions

The experimental strategy was conceived by J.-T.C. and P.Z. and further developed by C.-K.C.D. and J.R‥ X-ray crystallography and enzymatic assays were performed by J.R. and analyzed by J.R. and P.Z. Mass spectrometry measurements were done by Z.G. Measurements of the enzymatic activity and NADP(H) concentration in the cell lysates were done by C.-K.C.D., J.W., and J.R‥ Cell culture and transcriptome profiling experiments were done by C.-K.C.D. with assistance from J.W., T.S., P.-H.C., E.X., S.T., and J.A. RNAseq and microarray data were analyzed by K.-Y.C. and C.-K.C.D. C.-K.C.D., J.R., J.-T.C. and P.Z. wrote the manuscript with input from all co-authors.

## Data deposition

The coordinate of the hMesh1 D66K-NADPH complex has been deposited at the PDB databank, with the accession number of 5VXA. The RNA-Seq and microarray data have been deposited into NCBI GEO with accession number: GSE114282 (RNAseq) and GSE114128 (cDNA microarray for siMESH1 and siATF4).

## Competing financial interests statement

The authors declare no conflicts of interests.

